# Insulin potentiates JAK/STAT signaling to broadly inhibit flavivirus replication in insect vectors

**DOI:** 10.1101/701714

**Authors:** Laura R.H. Ahlers, Chasity E. Trammell, Grace F. Carrell, Sophie Mackinnon, Brandi K. Torrevillas, Clement Y. Chow, Shirley Luckhart, Alan G. Goodman

**Author notes:** Correspondence; @GoodmanLabWSU.

## Abstract

The World Health Organization estimates that over half of the world’s population is at risk for vector-borne diseases, such as those caused by arboviral infection. Because many arboviruses are mosquito-borne, investigation of the insect immune response will help identify targets that could reduce the spread of these viruses by the mosquito. In this study, we used a genetic screening approach to identify *insulin-like receptor* as a novel component of the immune response to arboviral infection. We determined that vertebrate insulin reduces West Nile virus (WNV) replication in *Drosophila melanogaster* as well as WNV, Zika, and dengue virus titers in mosquito cells. Mechanistically, we showed that insulin signaling activates the JAK/STAT, but not RNAi, pathway to control infection. Finally, we validated that insulin priming of adult female *Culex* mosquitoes through a blood meal reduces WNV infection, demonstrating an essential role for insulin signaling in insect antiviral responses to emerging human pathogens.

## INTRODUCTION

The World Health Organization (WHO) estimates that over half of the world’s population is at risk for contracting a vector-borne disease. Mosquitoes are the most common insect vector, and they transmit the flaviviruses West Nile virus (WNV), dengue virus (DENV), and Zika virus (ZIKV). *Culex spp.* are important mosquito vectors and were the most frequently infected with WNV in the U.S. in 2017 (ArboNET, 2018). Since 1999, WNV infection has caused disease in the 48 continental states, frequently resulting in mild, febrile symptoms. In more extreme cases, WNV causes neuroinvasive infection with severe complications and long-term patient consequences, including paralysis (Petersen et al., 2013). In 2017, the Centers for Disease Control (CDC) reported 2,097 cases of human WNV infection in the United States with 68% being classified as neuroinvasive (ArboNET, 2018). DENV affects 390 million people worldwide each year (Bhatt et al., 2013). ZIKV, another arbovirus with significant health impacts, was originally confined to small outbreaks, but expanded rapidly to over 30 countries between 2015 and 2016 (Tham et al., 2018). Significantly, there are currently no post-exposure therapeutics or effective vaccines against WNV, DENV, or ZIKV, indicating an unmet need in public health.

Insects use broadly antiviral signaling pathways to respond to virus infection, most notably the RNA interference (RNAi) response and the JAK/STAT pathways. *Drosophila melanogaster* activates RNAi as an innate immune response to DNA and RNA viruses (Bronkhorst et al., 2012; van Rij et al., 2006), and specifically to WNV (Chotkowski et al., 2008) and DENV (Mukherjee and Hanley, 2010). Subsequently it was shown that the RNAi pathway responds to WNV subtype Kunjin virus (WNV-Kun) infection in *Cx. quinquefasciatus* adult females (Paradkar et al., 2014). Studies using *D. melanogaster* found that RNAi pathway components communicate with non-canonical signaling proteins. For example, Deddouche, *et al*. demonstrated that the gene *vago* is induced by *Drosophila* C virus (DCV) infection in a Dicer-2 (Dcr-2)-dependent manner (Deddouche et al., 2008), and the transcription factor FoxO binds to the promoter regions of the RNAi components Dcr-2, Argonaute 1 (AGO1), and AGO2 (Spellberg and Marr, 2015) during arboviral infection.

Notably, WNV, DENV (Schnettler et al., 2012), and WNV-Kun (Moon et al., 2015) generate subgenomic flavivirus RNA (sfRNA) to antagonize the RNAi response and enhance virus survival. In addition, *D. melanogaster* uses the JAK/STAT pathway to respond to many RNA viruses, including DCV, cricket paralysis virus (CrPV), and Sindbis virus (SINV) (Kemp et al., 2013), as well as the DNA virus invertebrate iridescent virus 6 (IIV-6) (West and Silverman, 2018). The JAK/STAT pathway is activated when secreted cytokines of the unpaired (upd) family engage the cell surface receptor domeless (dome) (Brown et al., 2001; Harrison et al., 1998). When activated, dome signals through the tyrosine kinase hopscotch (hop) (Binari and Perrimon, 1994) leading to phosphorylation of the transcription factor Stat92E (Yan et al., 1996). Stat92E controls the induction of genes that regulate cell proliferation, organismal growth, stem cell renewal, development, and immunity (Arbouzova and Zeidler, 2006). During viral infection in *D. melanogaster*, Stat92E induces *vir-1* (Dostert et al., 2005) and *Turandot M* (*TotM*) (Kemp et al., 2013), whose protein products have antiviral activity. *Aedes aegypti* uses the JAK/STAT pathway to respond to the flaviviruses WNV, DENV, and yellow fever virus (YFV) (Colpitts et al., 2011) and JAK/STAT signaling is responsive to WNV infection in *Cx. quinquefasciatus* cells (Paradkar et al., 2012). While it is known that the RNAi and JAK/STAT responses are important for controlling arboviral infection in insects, it is not known whether these pathways communicate with each other or if there are additional, but as yet unidentified, signaling components that contribute to antiviral immunity.

Genetic variation in the human population contributes to disease progression for WNV (Bigham et al., 2011; Rios et al., 2010), DENV (Pabalan et al., 2017; Xavier-Carvalho et al., 2017), and ZIKV (Rossi et al., 2018). WNV causes symptoms in only 20% of infected individuals, (Hadler et al., 2014) with two-thirds of those cases becoming neuroinvasive (ArboNET, 2018), DENV manifests in approximately 25% of individuals (Bhatt et al., 2013), and ZIKV is symptomatic in 18% of individuals (Duffy et al., 2009). Therefore, there is power in using genetic screens to uncover potential risk factors or gene mutations that contribute to flavivirus infection. For example, a genetic screen in humans revealed a single-nucleotide polymorphism (SNP) in *OAS1* that is associated with symptomatic WNV infection (Bigham et al., 2011). Rios, *et al* also identified mutations in equine *OAS1* that are associated with WNV disease (Rios et al., 2007, 2010). Similarly, genetic screens can be used to identify pathways that control innate immune responses or viral load in the arthropod vector (Kingsolver et al., 2013). Considering the remarkable genetic diversity of insect viruses (Li et al., 2015; Webster et al., 2015), the identification of novel antiviral pathways in insect vectors is important in understanding viral escape mechanisms or entry points and the development of viral control methods (Marques and Imler, 2016)

In this study, we used the *Drosophila* Genetics Reference Panel (DGRP), a fully sequenced, inbred panel of fly lines derived from a natural population (Mackay et al., 2012), to perform a pathway-unbiased screen for natural genetic variants associated with WNV-Kun infection. Through our screen, we found that the insulin receptor (*InR*) was necessary for host survival to WNV-Kun. We also determined that priming insect cells with vertebrate insulin activates insulin signaling through Akt and ERK and induces prolonged transcription of JAK/STAT-mediated antiviral genes. The effect of insulin priming was antiviral in both flies and mosquitoes and antiviral to other flaviviruses, namely ZIKV and DENV. This work identifies insulin signaling as a novel component of insect immunity to arboviral infection and demonstrates that insulin signaling works cooperatively with known antiviral immune pathways for host protection.

## RESULTS

We used the *Drosophila* Genetic Reference Panel (DGRP) (Mackay et al., 2012) and a corresponding genome-wide association study (GWAS) (Chow et al., 2013, 2016; Lavoy et al., 2018) to determine how natural genetic variation in flies can lead to inter-individual differences in survival during viral infection. The DGRP has been used to identify novel genes that respond to endoplasmic reticulum stress (Chow et al., 2013), bacterial infection (Bou Sleiman et al., 2015; Howick and Lazzaro, 2017; Wang et al., 2017), and to lead toxicity (Zhou et al., 2016). Unbiased GWAS is informative in determining the genes associated with a particular phenotype and can identify novel genes connected with the phenotype of interest (Stranger et al., 2011). We performed our screen using the naturally attenuated WNV subtype Kunjin virus (WNV-Kun), which has high sequence identity to the Lineage I WNV-New York strain (Lanciotti et al., 2002) and can infect *D. melanogaster* (Yasunaga et al., 2014). WNV-Kun can be used in arthropod containment level 2 (ACL2) facilities (Hackett and Cherry, 2018; U.S. Department of Health and Human Services, 2009).

We infected female flies from 94 of the DGRP lines with 104 plaque-forming units (PFU) WNV-Kun by intrathoracic injection and determined their mortality rates as compared to flies treated with buffer only (mock infection) (Fig. 1A). In this experiment, we used fly lines that were free of the endosymbiont *Wolbachia* since infection with this organism can protect against RNA virus infection (Teixeira et al., 2008). We monitored survival daily for 30 days and calculated the hazard ratio as a metric of survival (Fig. 1B). We then used the log of the hazard ratio as the quantitative phenotype for the GWAS (Table S1), as described (Chow et al., 2013), and identified a set of genome-wide suggestive variants (nominal *P* < 5×10-5, Table S2), including *InR*. Next, we performed gene set enrichment analysis (GSEA) (Fig. 1C) to identify groups of gene variants that were functionally enriched. Input for GSEA consisted of the gene and P-value associated with each variant from the entire dataset. Given a defined set of genes annotated with a certain gene ontology (GO) function, GSEA determines how members of a GO category are distributed throughout the list of genes ranked by P-value. GO categories enriched at the top of the list functionally describe the phenotype of the gene set (Subramanian et al., 2005). In the GO category regulation of cell proliferation (GO: 0008284), we identified genes that were previously shown to play a role in fly immunity to RNA virus infection via the JAK/STAT pathway (orange boxes), namely *Stat92E*, *dome*, *upd3*, and *caudal* (*cad*) (Zhou and Agaisse, 2012). In addition to *InR*, we identified a number of insulin-like peptides (*ilps*) that were part of the functionally enriched GO categories female mating behavior (GO: 0060180) and hormone activity (GO: 0005179) (blue boxes). Functional enrichment of these *ilps* supported a role for insulin signaling in the antiviral immune response.

**Figure 1:**
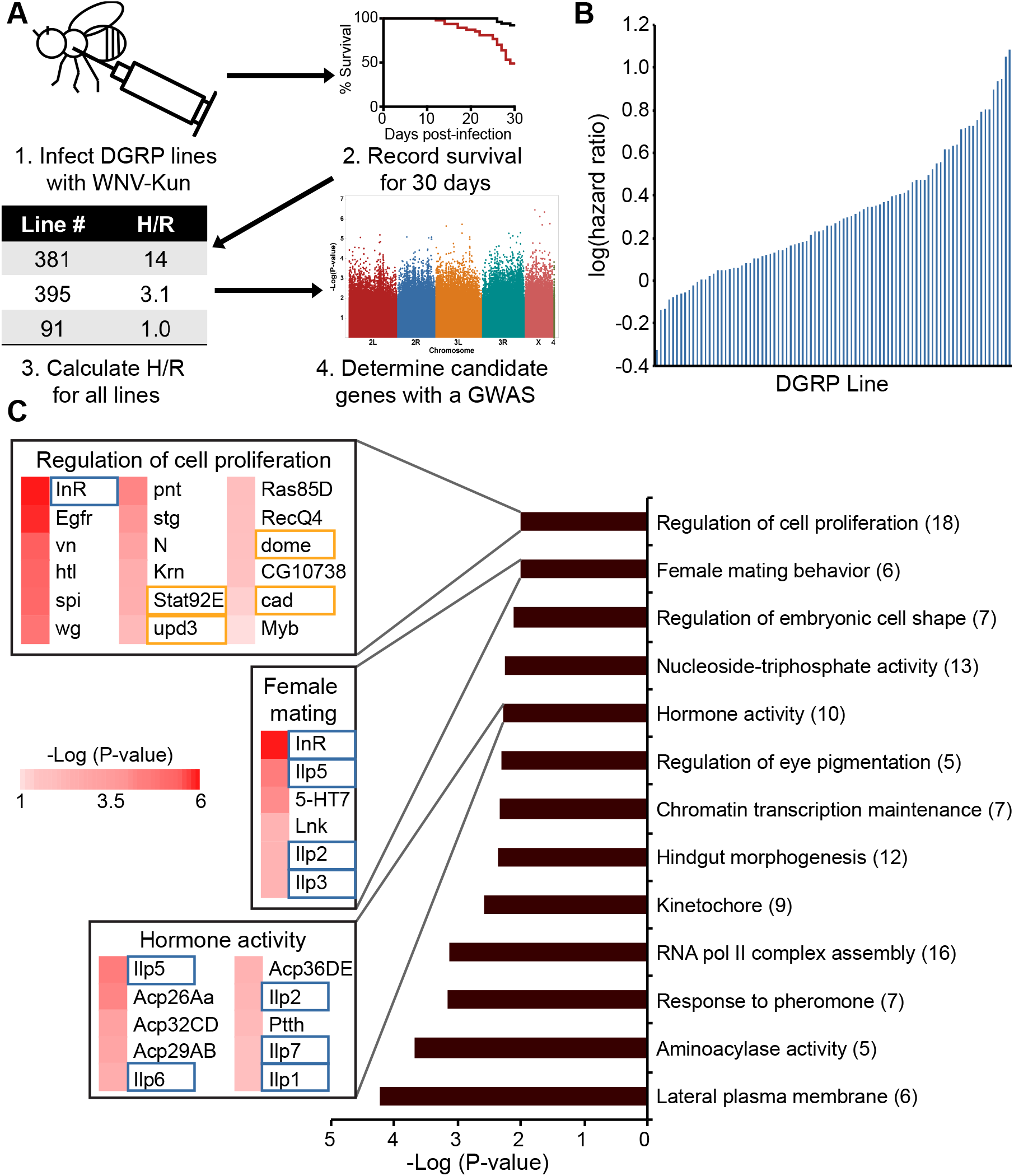
A genetic screen of *D. melanogaster* identified candidate genes involved in the *D. melanogaster* response to WNV-Kun. (A) Schematic of the screen and downstream analysis. (B) Survival of each DGRP line, measured by log(hazard ratio). (C) Gene set enrichment analysis (GSEA) using genes for all variants and their associated P-values from the GWAS. Heatmap data represent GWAS variant P-values while the bar graph indicates the GSEA P-value and the number of genes enriched for each GO category in parentheses. Genes indicated in orange boxes are components of the JAK/STAT pathway, and genes indicated by blue boxes are components of the insulin response pathway.

We selected *upd3*, *Stat92E, hop*, and *InR* for validation, but also included *tak1* (TGFβ-activated kinase 1) and *egfr* (epidermal growth factor receptor) based on previously identified roles for these genes in RNA virus infection (Xia et al., 2017). *Vir-1* is induced downstream of the JAK/STAT pathway (Kemp et al., 2013) and was used as a surrogate to validate this pathway. Fly lines with these genes deleted or knocked-down by RNAi were infected with WNV-Kun. Because *upd2* and *upd3* have redundant roles (Kemp et al., 2013), we selected a line with mutations in both *upd2* and *upd3* for these analyses. Mortality rates for each genotype during WNV-Kun infection were compared to mock infected (buffer only) controls. As expected, JAK/STAT pathway mutants were susceptible to WNV-Kun infection based on increased mortality (P-value < 0.05 and hazard ratio >1) in *upd^Δ2^upd^Δ3^* and *hop* mutants, while the control line *y^1^w^1^* did not display significant mortality (Fig. 2A-C). Viral titers in *Stat92E* and *vir-1* mutants were also increased relative to controls (Fig. 2D). Our data confirmed that knockdown of *InR* or *tak1* increased susceptibility to WNV-Kun infection (Fig. 2E-F and Fig. S1A), whereas *egfr* knockdown, in contrast, did not display significant mortality (Fig. S1B). Thus, we validated DGRP candidate genes *InR* and *tak1* as well as JAK/STAT pathway genes previously associated with RNA virus infection.

**Figure 2:**
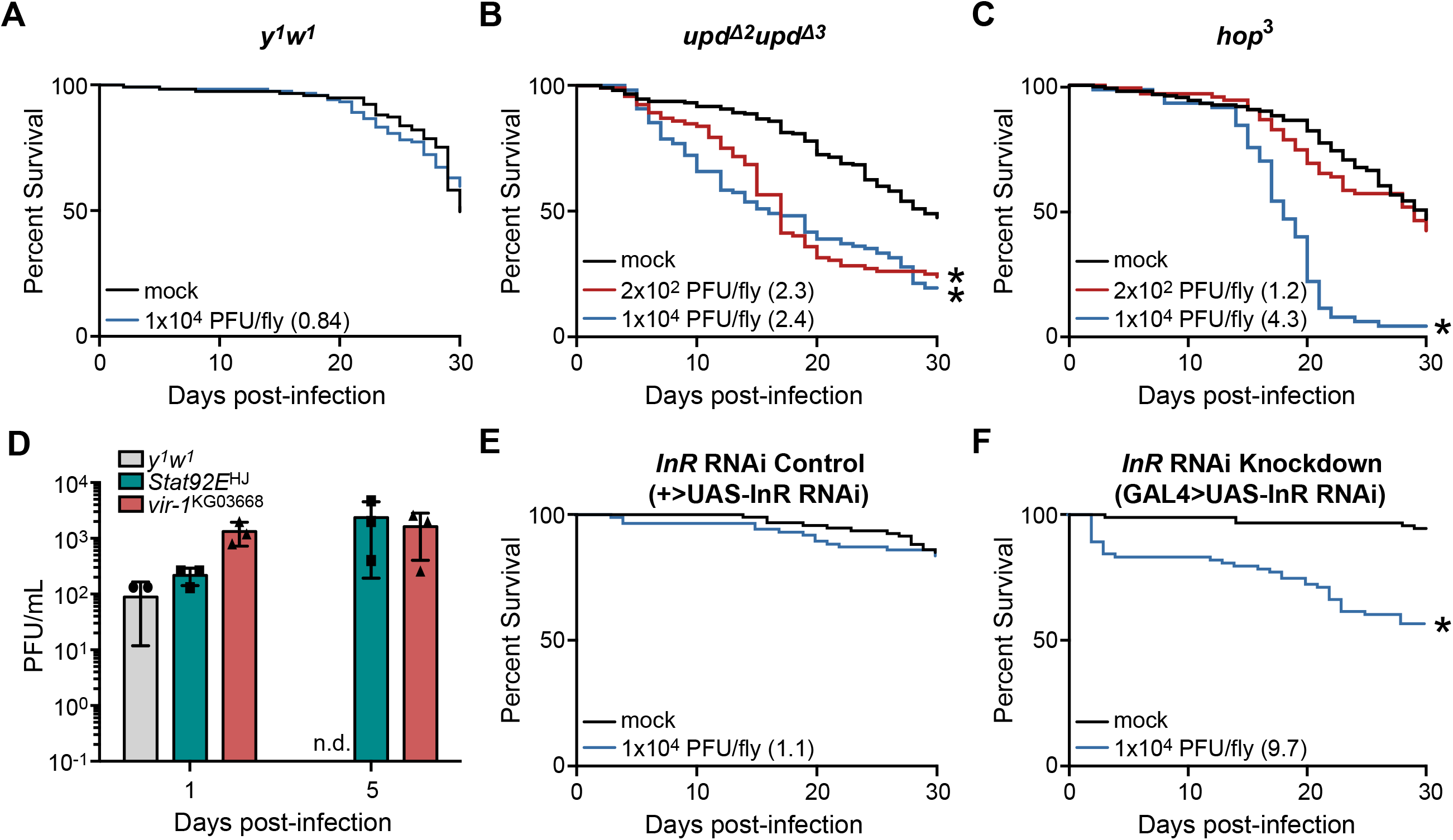
Components of the JAK/STAT and insulin response pathways are necessary for *D. melanogaster* survival to WNV-Kun infection. (A-C) Mutants in the genes (B) *upd2* and *upd3* and (C) *hop* are susceptible to WNV-Kun infection compared to the (A) *y^1^w^1^* isotype control. (D) WNV-Kun titer is higher in *Stat92E* and *vir-1* mutant flies compared to the *y^1^w^1^* isotype control. (E-F) *InR* knockdown flies are susceptible to WNV-Kun compared to the sibling controls. Hazard ratio for each infection group is indicated in parenthesis, and statistical significance from the mock infection group is indicated with an asterisk. (Log-rank test; *p < 0.05). Each survival curve represents two independent experiments of >40 flies that were combined for a final survival curve. Error bars represent SDs. Samples where virus was not detected are indicated by “n.d

Based on validation of *InR* and the identification of *ilps* using GO analysis, we tested the effects of insulin on the activation of Akt and WNV-Kun replication in *D. melanogaster* S2 cells and on virus replication in adult flies. The insulin signaling pathway in *D. melanogaster* is initiated by insulin-like peptide (ilp) binding to the insulin receptor (InR), which induces a phosphorylation cascade through the signaling protein chico, with subsequent bifurcation through two pathways, one of which is dependent on activation of PI3K (phosphoinositide 3-kinase), PDK1 (pyruvate dehydrogenase kinase 1), and Akt (Puig et al., 2003) and a second that is dependent on the activation of MAPK/ERK with feedback between the two branches (reviewed in Luckhart and Riehle, 2007). In *D. melanogaster* S2 cells, we observed robust Akt phosphorylation following 8 hours of 1.7 μM insulin treatment (Fig. 3A). Moreover, we observed reduced transcript levels of *InR* during exogenous insulin treatment, as measured by qRT-PCR (Fig. S2). This was expected, as insulin-induced activation of Akt-mediated FoxO phosphorylation results in cytoplasmic retention of FoxO and a concomitant loss of FoxO-dependent *InR* induction (Puig et al., 2003). After 24 hours of continuous insulin treatment prior to infection, WNV-Kun replication was also significantly reduced in S2 cells (Fig. 3B), suggesting that insulin-mediated Akt activation stimulated an antiviral response. We tested insulin control of virus replication *in vivo* using WNV-Kun-infected OregonR flies raised on food containing 10 μM insulin. As in S2 cells, viral titer was significantly reduced in insulin-treated flies relative to control flies by 10 days post-infection (Fig. 3C).

**Figure 3:**
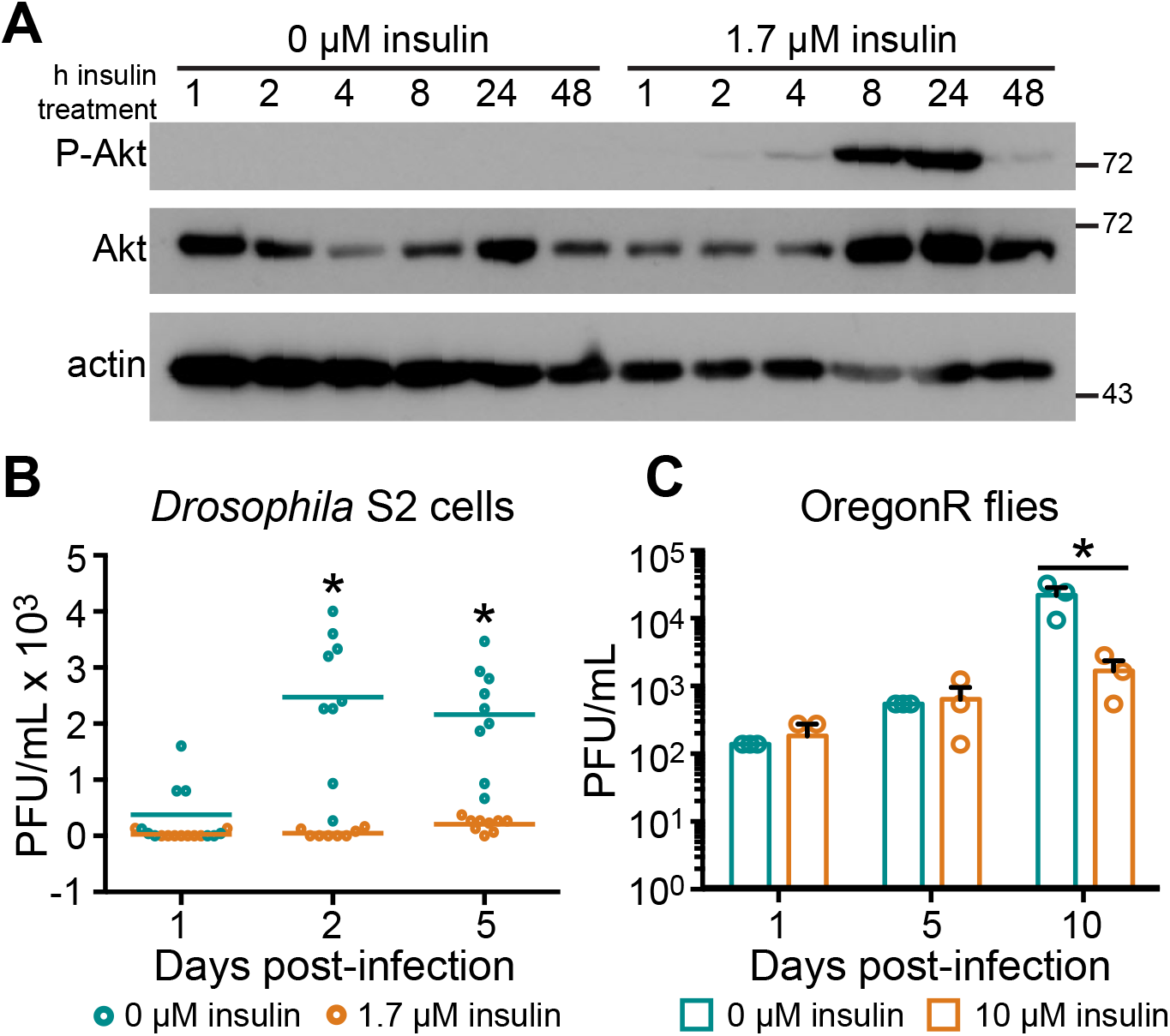
The *D. melanogaster* insulin response pathway is activated by vertebrate insulin, which is antiviral to WNV-Kun. (A) Akt is phosphorylated and activated in *D. melanogaster* S2 cells when treated with bovine insulin. (B) WNV-Kun titer is reduced in S2 cells primed with 1.7 μM bovine insulin for 24 hours prior to infection (MOI 0.01 PFU/cell). (C) WNV-Kun titer is reduced in *D. melanogaster* on a diet of 10 μM bovine insulin. (Unpaired t-test; *p < 0.05). Results are representative of duplicate experiments. Error bars represent SDs.

We next investigated signaling crosstalk during insulin-mediated restriction of WNV-Kun replication with particular focus on JAK/STAT, RNAi, and ERK signaling. We determined that insulin treatment induced prolonged transcription of JAK/STAT components *upd2, upd3* and *TotM* (Fig. 4A-C), while significant *vir-1* induction was observed during insulin treatment only at 8 hours post-infection (Fig. 4D). In all cases, these effects were independent of virus infection (Fig. 4A-D). The RNAi components *AGO1, AGO2* and *Dcr-2* were induced at early times post-infection during insulin treatment (Fig. 4E-G). However, by 24 hours post-infection, *AGO1* induction was significantly reduced during insulin treatment in both control and virus-infected cells (Fig. 4E), *AGO2* induction was not observed in either cell treatment (Fig. 4F), and *Dcr2* induction persisted only in control cells (Fig. 4G). Together, these results suggested that insulin exposure prolongs the induction of JAK/STAT components, but not RNAi components, indicating that insulin sustains JAK/STAT signaling for antiviral activity.

**Figure 4:**
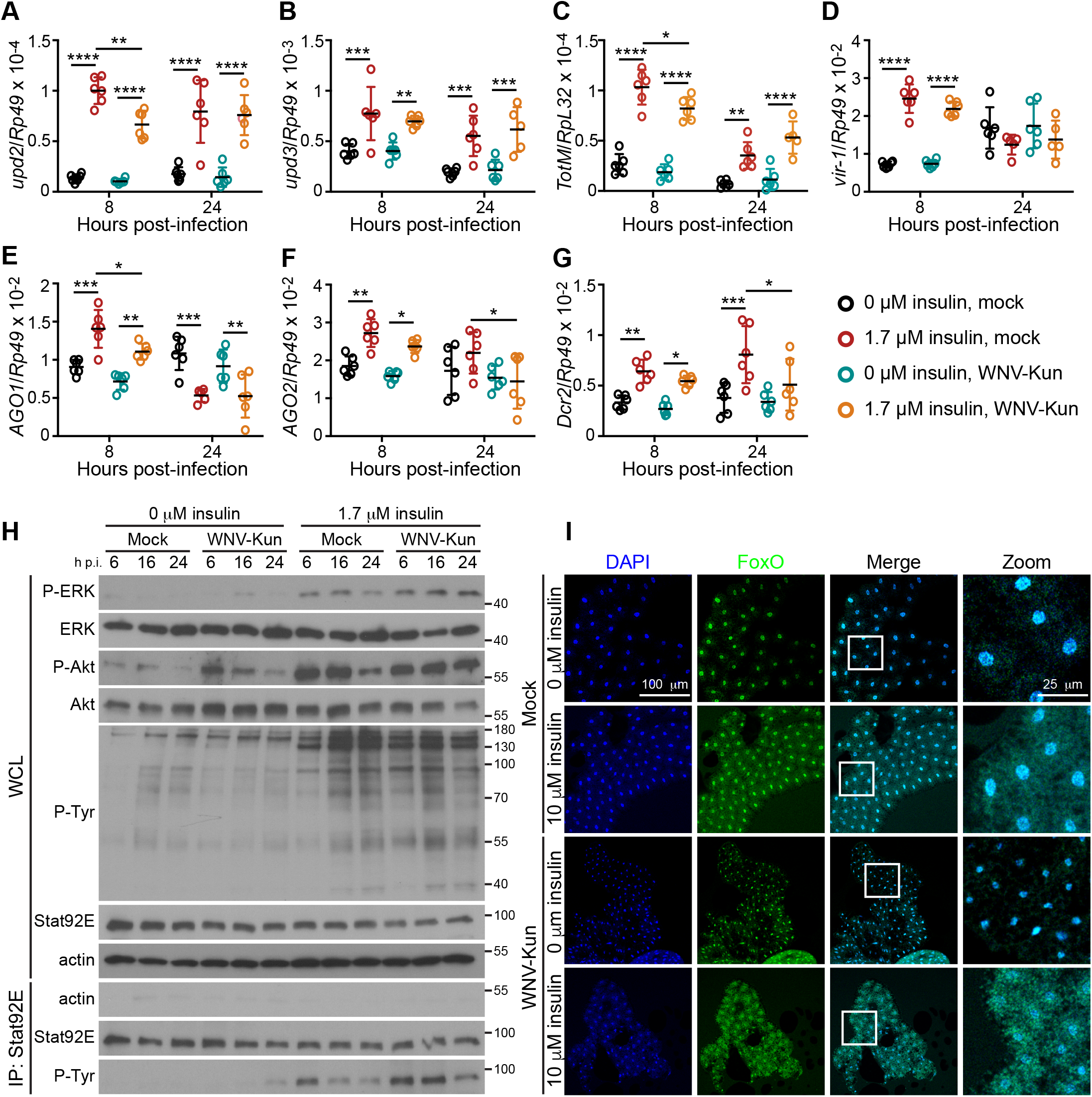
Insulin priming activates antiviral pathways in *D. melanogaster*. Induction of genes within the (A-D) JAK/STAT pathway and the (E-G) RNAi pathway were measured by qRT-PCR following priming of *D. melanogaster* S2 cells with 1.7 μM insulin and mock or WNV-Kun infection. (H) Levels of ERK, Akt, and Stat92E phosphorylation were measured by western blot following insulin treatment and mock or WNV-Kun infection of S2 cells. (I) FoxO localization in the larval fat body was determined by confocal microscopy using 3rd instar larvae on a diet of 10 μM bovine insulin and mock- or WNV-Kun-infected for 4 hours. (ANOVA with correction for multiple comparisons; *p < 0.05; **p<0.01; ***p<0.001; ****p<0.0001). Results are representative of duplicate experiments. Error bars represent SDs.

To validate associations between insulin and JAK/STAT signaling, we examined temporal activation (phosphorylation) of ERK, Akt, and Stat92E in control and WNV-Kun-infected S2 cells with and without insulin treatment. ERK was phosphorylated during insulin treatment as shown previously (Xu et al., 2013) and this phosphorylation was elevated during WNV-Kun infection relative to control (mock; Fig. 4H**, top row**). Akt phosphorylation was induced by WNV-Kun infection and this was enhanced by insulin treatment relative to control (Fig. 4H**, 3rd row from top**). Low levels of Akt activation were observed during infection in the absence of insulin, which was expected based on previous results showing that WNV activates Akt (Shives et al., 2014). In support of a role for JAK/STAT signaling during insulin treatment, we immunoprecipitated Stat92E from whole S2 cell lysate to probe for tyrosine phosphorylation indicative of Stat92E activation. Similar to ERK, Stat92E was activated during WNV-Kun infection and phosphorylation was enhanced relative to controls (mock) by insulin treatment (Fig. 4H**, bottom row**).

Given that FoxO activity induces expression of RNAi components that are antiviral during CrPV infection (Spellberg and Marr, 2015), we sought to examine the role of FoxO during insulin treatment and WNV-Kun infection. For this purpose, a cohort of *D. melanogaster* engineered with a FoxO-GFP reporter was reared on food containing 10 μM insulin and infected as third instar larvae with WNV-Kun. Given that the fat body is an important immune organ during viral infection in flies (Lautié-Harivel and Thomas-Orillard, 1990), we dissected this tissue to visualize FoxO-GFP localization by confocal microscopy from treated and control larvae. As expected, insulin treatment increased cytosolic FoxO-GFP and this effect was independent of WNV-Kun infection (Fig. 4I). Hence, insulin treatment results in a loss of FoxO-dependent transcription, consistent with the loss of RNAi gene product induction during prolonged insulin treatment and infection (Fig. 4E-G). In this context, ERK phosphorylation would likely lead to the activation of JAK/STAT signaling and target genes for enhanced antiviral activity during insulin treatment.

Since mosquitoes are the natural vectors for WNV, we sought to determine whether our findings in *D. melanogaster* would translate to mosquitoes. To this end, we determined that insulin treatment of *Cx. quinquefasciatus* Hsu cells and *Ae. albopictus* C6/36 cells activated Akt (Fig. 5A-B) and reduced titers of WNV-Kun relative to control, untreated cells (Fig. 5C-D). This antiviral effect was not limited to WNV-Kun in that insulin treatment reduced the titers of two additional flaviviruses, ZIKV and DENV, in C6/36 cells (Fig. 5E-F). Notably, C6/36 cells lack an RNAi response (Brackney et al., 2010), demonstrating that an RNAi response is not required for insulin-mediated antiviral activity. Finally, WNV-Kun titer was increased significantly in *Ae. albopictus* C6/36 cells treated with 10 μM MEK/ERK inhibitor U0126 (Fig. S3) and this treatment reversed the effects of insulin on virus replication (Fig. S3), affirming that virus-and insulin-induced ERK activation (Fig. 4H**, top row**) are functionally important in the regulation of virus replication in mosquito cells (Fig. S3).

**Figure 5:**
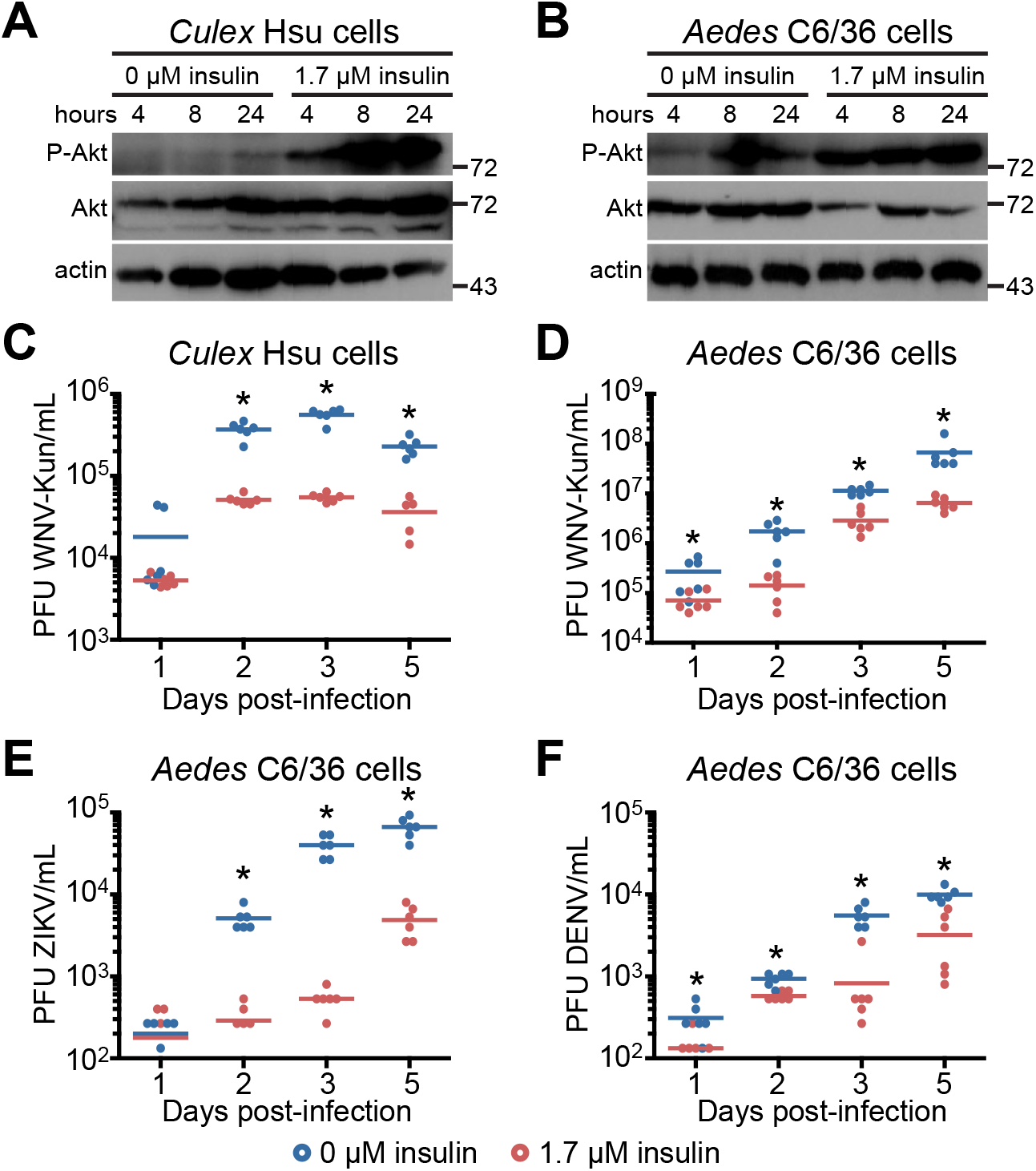
Insulin priming reduces flavivirus titer in *Cx. quinquefasciatus* and *Ae. albopictus* cells. Akt is phosphorylated and activated by insulin priming in (A) *Cx. quinquefasciatus* Hsu cells and (B) *Ae. albopictus* C6/36 cells. (C-D) Insulin priming reduces WNV-Kun titer in Hsu and C6/36 cells, and (E) Zika virus and (F) dengue virus titer in C6/36 cells (MOI 0.01 PFU/cell). (Unpaired t-test; *p < 0.05). Results are representative of duplicate experiments. Error bars represent SDs.

Lastly, we tested the effects of insulin treatment on WNV-Kun replication in *Cx. quinquefasciatus* adult females. Age-matched 6-9 day old adult female mosquitoes were infected with WNV-Kun via blood meal in the presence or absence of insulin and collected at 1, 5, and 10 days post-infection (Fig. 6A). Mosquitoes that did not blood feed were excluded from subsequent analyses. We selected 170 pM bovine insulin, as this dose is within the physiological range in humans (Darby et al., 2001) and activates insulin signaling and alters *Plasmodium falciparum* development in *Anopheles stephensi* (Pakpour et al., 2012). Similar to our results in *D. melanogaster,* we observed that insulin treatment lead to decreased *R2D2* induction in insulin-fed control mosquitoes (Fig. 6B), suggesting a reduced RNAi response in the context of increased *STAT* expression in insulin-fed infected mosquitoes (Fig. 6C). Further, WNV-Kun *env* and *NS5* gene expression levels were reduced in insulin-fed mosquitoes by 10 days post-infection (Fig. 6D-E). While insulin-induced *STAT* expression was significant at 1 day post-infection (Fig. 6C), markers of virus infection were not significantly reduced until 10 days post-infection (Fig. 6D-E). Intriguingly, numerous reports of WNV infection in mammalian cells indicate that WNV non-structural proteins, including NS5 for which expression increases over time in both control and insulin-treated mosquitoes (Fig. 6E), can interfere with JAK/STAT signaling (Guo et al., 2005; Laurent-Rolle et al., 2010; Muñoz-Jordán et al., 2005), suggesting an explanation for the delay in observed effects of insulin on virus infection. Moreover, due to a point mutation in NS5, WNV-Kun displays reduced STAT antagonism (Laurent-Rolle et al., 2010), which allows for a more robust JAK/STAT host response during infections with this strain compared to virulent strains of WNV. Collectively, these results suggest that vertebrate insulin ingested with the blood meal limits virus infection in the vector, perhaps via a STAT-mediated immune response that enables the vector to survive infection and transmit the virus to another host.

**Figure 6:**
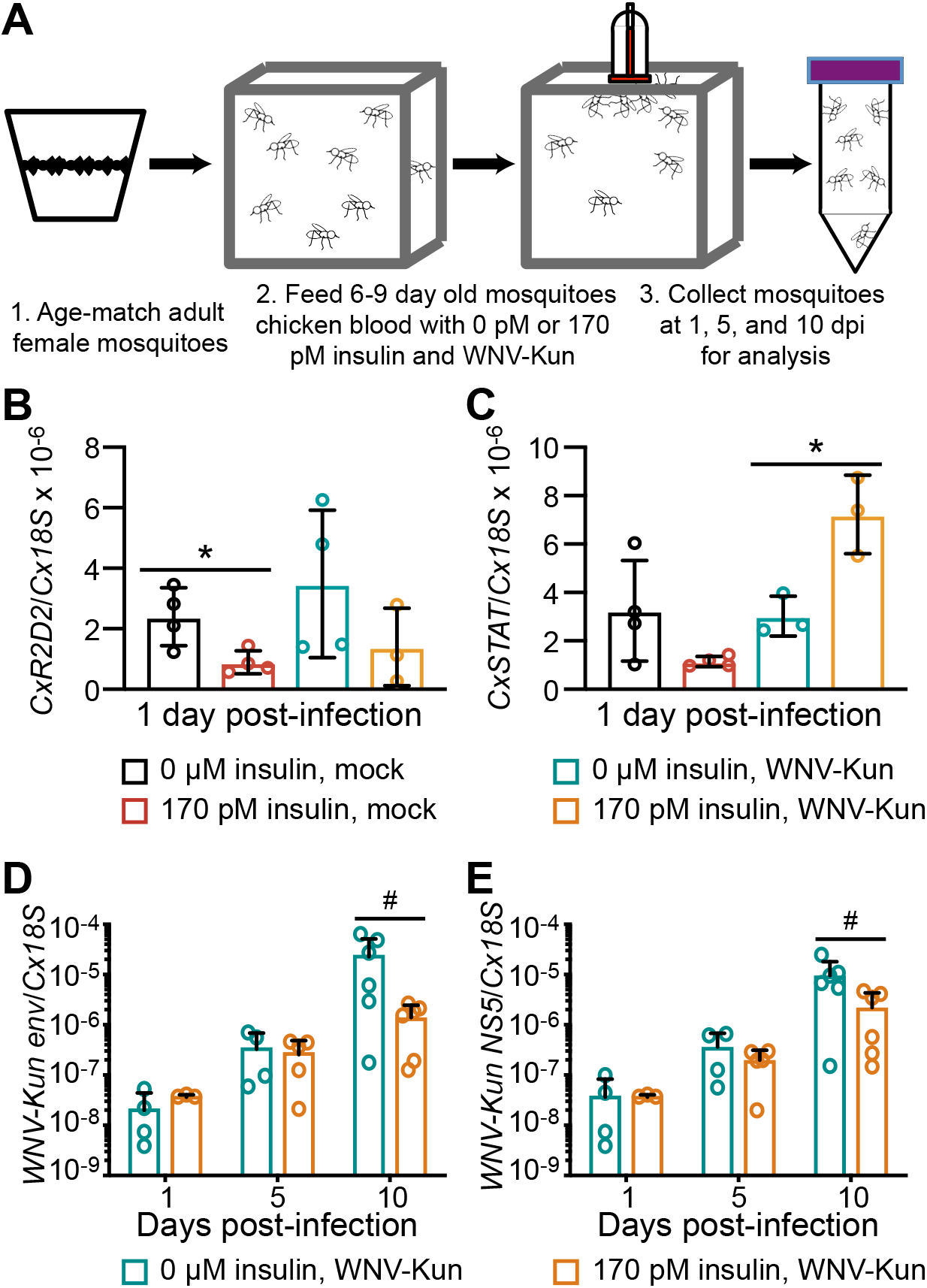
Insulin activates the antiviral response in adult female *Cx. quinquefasciatus*. (A) Schematic illustrating process of age-matching pupae into adults, feeding female mosquitoes a blood meal of chicken blood with or without insulin or WNV-Kun, and collecting adults for analysis post-infection. (B-E) Induction of (B) *CxR2D2,* (C) *CxSTAT*, (D) WNV-Kun *envelope* (*env*), and (E) *NS5* genes were measured by qRT-PCR following blood feeding of *Cx. quinquefasciatus*. (Unpaired t-test; *p < 0.05 or Mann-Whitney test; #p < 0.05). Results are representative of duplicate experiments. Error bars represent SDs.

## DISCUSSION

In the work presented here, we used the DGRP for the first time to investigate an antimicrobial host response to flaviviral infection and identified *InR*, along with other known antiviral response mediators, as components of the *D. melanogaster* response to WNV-Kun infection. We demonstrated that vertebrate insulin activates insulin and JAK/STAT signaling for a net antiviral effect in *D. melanogaster* cells. Moreover, an analogous antiviral effect is detectable in *Cx. quinquefasciatus* and *Ae. albopictus* cells, as well as *Cx. quinquefasciatus* females, and this insulin-dependent response is broadly antiviral to other flaviviruses, including DENV and ZIKV. Collectively, our data suggest that insulin activates known immune response pathways *in vitro* and *in vivo* for overall host restriction of flavivirus infection.

Mechanistically, we show that *InR* mediates its broad anti-flaviviral activity by selectively potentiating the JAK/STAT but not RNAi response. We show that ERK is activated in the context of FoxO inactivation and cytosolic localization, and we hypothesize that ERK links insulin signaling to the JAK/STAT pathway (Fig. 7). Among elements related to JAK/STAT signaling in flies, upd2 controls ilp secretion by the fat body during the fed state (Rajan and Perrimon, 2012). The JAK/STAT-dependent genes *vir-1* and *TotM* are induced during RNA virus infection in *D. melanogaster,* with rapid-kill viruses (< 10 days) inducing a *vir-1* response and slow-kill viruses inducing a *TotM* response. WNV-Kun typically kills flies slowly, substantiating the prolonged *TotM* induction (Fig. 4C) that we observed. Given that MEKK1 signaling can contribute to *TotM* induction (Brun et al., 2006), ERK activation could prolong *TotM* induction during insulin priming to enhance an overall antiviral effect through JAK/STAT. In addition, we observed that ERK activation by insulin was induced concurrently with Stat92E phosphorylation (Fig. 4H) and directly regulated virus replication (Fig. S3), while FoxO activation and expression levels of RNAi components were reduced, suggesting that insulin activates the JAK/STAT pathway and ERK to control WNV-Kun replication. Interestingly, we observed that the antiviral RNAi response was not reliant on insulin signaling. This may be due, in part, to the fact that insulin signaling and a nutrient-rich environment lead to FoxO phosphorylation and reduced induction of FoxO target genes, such as *InR* and those that encode RNAi machinery (Puig et al., 2003; Spellberg and Marr, 2015). While these conditions would lead to increased growth and lifespan regulation in *D. melanogaster* (Giannakou et al., 2004; Hwangbo et al., 2004) and mosquitoes (Arik et al., 2015; Kang et al., 2008), in the context of viral infection, they could be detrimental due to decreased antiviral RNAi activity. However, our results support a model in which insulin induced JAK/STAT signaling compensates for the loss of RNAi activity. Importantly, we observed an insulin-mediated antiviral phenotype in *Ae. albopictus* C6/36 cells, which lack an RNAi response (Brackney et al., 2010), providing a natural experiment in support of our inference that the insulin-dependent mosquito antiviral response is not mediated through RNAi.

**Figure 7:**
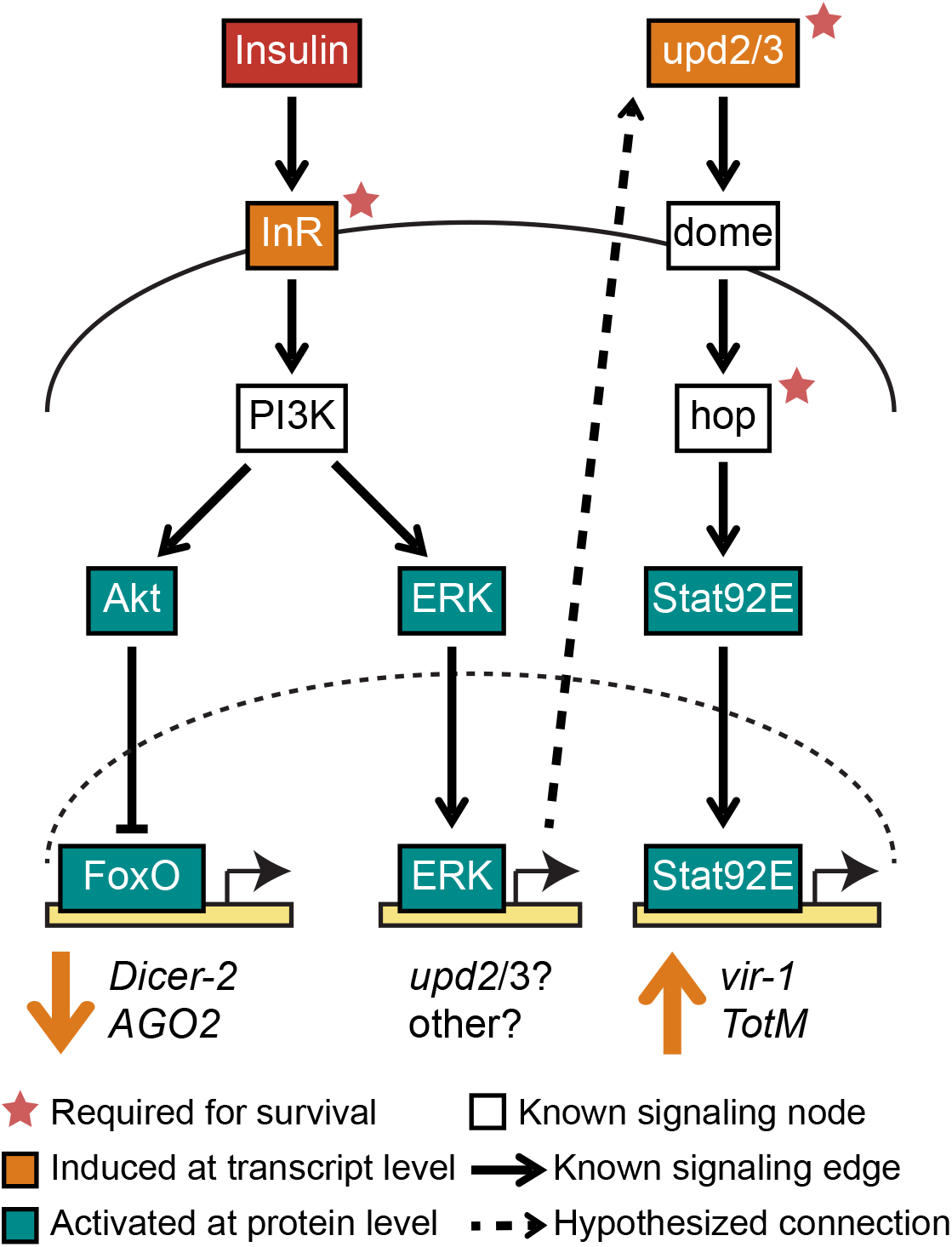
Schematic of immune signaling during insulin priming in insects. Insulin (red box) binds to the insulin receptor (InR), activating a signaling cascade that inhibits FoxO-dependent transcription of RNAi components and reduces RNAi-dependent antiviral immunity. Proposed networked regulation of insulin and JAK/STAT signaling includes activation of ERK downstream of PI3K and increased expression of *upd2/3*, suggesting a control point for insulin-enhanced JAK/STAT signaling (dotted arrow). Signaling components denoted by pink stars were important for *D. melanogaster* survival during WNV-Kun infection. Components indicated by orange boxes were induced at the transcript level by insulin treatment, and those indicated by teal boxes were activated at the protein level. White boxes indicate known signaling nodes.

Prior to our studies, the insulin signaling pathway had not been specifically linked to host immunity to WNV. Intriguingly, however, diabetes mellitus has been shown to increase risk of WNV infection (Nash et al., 2001) and, in mouse models of the disease, leukocyte and T cell recruitment are decreased and WNV replication and neuroinvasiveness are increased in the absence of insulin (Kumar et al., 2012, 2014). These observations are mirrored in malaria parasite infection. Specifically, a higher percentage of *An. stephensi* become infected following feeding on *P. falciparum*-infected type 2 diabetic mice compared to mosquitoes that fed on control animals, suggesting that diabetic mice are more efficient at infecting mosquitoes (Pakpour et al., 2016). Indeed, insulin signaling has previously been implicated in immunity to parasites, bacteria, and viruses in both insects and mammals. In *D. melanogaster*, Musselman, *et al*. determined that insulin signaling is networked to peptidoglycan receptor signaling for an antibacterial immune response (Musselman et al., 2018). In *An. stephensi*, overexpression of activated Akt in the midgut blocked *P. falciparum* infection (Corby-Harris et al., 2010). In human cells, Akt signaling has been connected to the immune response to the flavivirus hepatitis C virus (HCV) (Aytug et al., 2003), and virus-induced disruption of insulin signaling was observed to increase HCV replication (Zhang et al., 2018). A variety of studies suggest that insulin signaling-dependent activation of ERK contributes to this antiviral effect. In particular, ERK is activated in response to DENV in human cells (Smith et al., 2014), by WNV in mouse cells (Scherbik and Brinton, 2010), and by vesicular stomatitis virus (VSV), SINV, and DCV in *D. melanogaster* cells and in adult flies (Xu et al., 2013). Furthermore, Xu, *et al*. showed that insulin- and ERK-dependent signaling restricts DCV and SINV infection in *D. melanogaster* and *Ae. aegypti* cells. Collectively, these studies and ours affirm that insulin signaling regulates innate immunity to a broad array of microbial pathogens.

Taken together, our data demonstrate that the *Cx. quinquefasciatus* response to insulin mirrors the *D. melanogaster* response in that JAK/STAT is induced by WNV-Kun infection, and is potentiated by insulin in the blood meal, perhaps through ERK signaling. Given this signaling framework and the conservation of this antiviral effect, we have leveraged fly genetics to efficiently identify and validate across species the downstream mechanisms that underlie insulin-dependent inhibition of viral replication. While we are cognizant of species-specific distinctions that likely exist across an estimated 260 million years of evolution between flies and mosquitoes (Arensburger et al., 2010), we have provided an example of using these two systems in parallel to speed the identification of novel gene targets involved in the regulation of flavivirus infection in vector mosquitoes. We suggest that we can combine this knowledge to engineer mosquito resistance to flavivirus infection and to leverage additional fly models to identify other vulnerable points in the disease cycle for control. In particular, given that diabetes is a risk factor for flaviviral disease (Guo et al., 2017; Mavrouli et al., 2018) and that ERK- and Akt-dependent pathways control viral replication in human cells as noted above, *D. melanogaster* models of diabetes (Inoue et al., 2018; Musselman and Kuhnlein, 2018) could extend our understanding of the mechanisms of increased risk for flavivirus infection and guide the identification of strategies to lessen human disease burden.

## ACKNOWLEDGEMENTS

We thank S. Best, A. Nicola, and R. Tesh for cells and viruses used in these experiments and the staff, particularly J. Huffman, at the Laboratory Animal Research Facility at the University of Idaho for assistance with mosquito husbandry. We thank A. Brown and J. Ahlers for expertise in analyzing bioinformatics data and S. Balachandran for critical reading of our manuscript.

This research was supported by NIH Grants R00 AI106963 and R21 AI128103 to A.G. Goodman, a NIH/NIGMS-funded pre-doctoral fellowship T32 GM008336 and a Poncin Fellowship to L.R.H. Ahlers, the Stanley L. Adler Research Fund, and the Mary V. Schindler Equine Research Endowment. S. Luckhart was supported by an NIH/NIAID R56 award (R56 AI118926). C.Y. Chow was supported by an NIH/NIGMS R35 award (R35 GM124780) and a Glenn Award from the Glenn Foundation for Medical Research. CYC is the University of Utah Mario R. Capecchi Endowed Chair in Genetics.

## AUTHOR CONTRIBUTIONS

Conceptualization, L.R.H.A. and A.G.G.; Methodology, L.R.H.A., C.E.T., G.F.C., B.K.T., C.Y.C., S.L.,and A.G.G.; Software, C.Y.C.; Validation, L.R.H.A., C.E.T., G.F.C., and S.M.; Investigation, L.R.H.A., C.E.T., G.F.C., S.M., B.K.T., C.Y.C., and A.G.G.; Resources, C.Y.C., S.L., and A.G.G.; Writing – Original Draft, L.R.H.A.; Writing – Review and Editing, C.E.T., G.F.C., S.M., C.Y.C., S.L., and A.G.G.; Visualization, L.R.H.A. and A.G.G.; Funding Acquisition, L.R.H.A., S.L., C.Y.C., and A.G.G.

## MATERIALS AND METHODS

### CONTACT FOR REAGENT AND RESOURCE SHARING

Further information and requests for resources and reagents should be directed to and will be fulfilled by the Lead Contact, Alan Goodman.

### EXPERIMENTAL MODEL AND SUBJECT DETAILS

#### Fly lines and genetics

A genetic screen was completed using the available fly lines from the DGRP (Mackay et al., 2012) that lack the endosymbiont *Wolbachia* (Bloomington *Drosophila* Stock Center). RNAi knockdown of *InR* was achieved by crossing a driver line for actin (actin5C-GAL4) with a fly line containing a cassette expressing the UAS for actin and dsRNA for the *InR* gene. Progeny flies were knocked-down for *InR* if they contained the actin5C-GAL4 driver, and the sibling control flies contained wild-type levels of *InR* if they carried the *CyO* balancer. Flies used in the study are listed in the Resources table. Flies were maintained on standard cornmeal food (Genesee Scientific #66-112) at 25 °C and 65% relative humidity. Bovine insulin (Sigma 10516) was added to food preparation for a final concentration of 10 μM, within the range described in (Xu et al., 2013).

#### Bioinformatics

GWAS analysis was completed as described in (Chow et al., 2013, 2016; Lavoy et al., 2018), using log(hazard ratio) as a metric of mortality (Chow et al., 2013). All single nucleotide polymorphism (SNP) coordinates are based on the *D. melanogaster* dm6 genome build.

Lines with less than 2% death in the mock-infected group were removed prior to analysis, as described (Chow et al., 2013). Gene Set Enrichment Analysis (GSEA) (Subramanian et al., 2005) was performed using all GWAS variant data and their associated P-values as previously described (Subramanian et al., 2005). The gene nearest to each variant was assigned the variant’s P-value and used as GSEA input. While traditional GO analysis uses a set of genes based on a P-value cutoff, GSEA examines the entire gene set (Dyer et al., 2008). A cut-off of *P* < 0.01 was used for the GO categories presented in Fig. 1C.

#### Cells and virus

Vero cells (ATCC, CRL-81) were kindly provided by A. Nicola and cultured at 37 °C/5% CO2 in DMEM (ThermoFisher 11965) supplemented with 10% FBS (Atlas BiologicalsFS-0500-A) and 1x antibiotic-antimycotic (ThermoFisher 15240062). *Aedes albopictus* C6/36 cells (ATCC, CRL-1660), which are free from persistent WNV, DENV, or ZIKV infection (Nag and Kramer, 2017; Nag et al., 2016), were cultured at 28 °C/5% CO2 in RPMI supplemented with 10% FBS (FisherScientific SH3007003HI), 0.15% sodium bicarbonate (ThermoFisher 25080), 1x non-essential amino acids (ThermoFisher 11140), 1x sodium pyruvate (ThermoFisher 11360), and 1x antibiotic-antimycotic. *Culex quinquefasciatus* Hsu cells (Hsu et al., 1970) were gifted by R. Tesh and cultured at 28 °C in L-15 medium (ThermoFisher 11415) supplemented with 10% FBS (FisherScientific SH3007003HI), 10% tryptose phosphate buffer (ThermoFisher 18050), and 1x antibiotic-antimycotic. Hsu cells have been shown to be free of densoviruses, which are found in other mosquito cell lines (O’Neill et al., 1995). S2 cells were cultured as described in (Ahlers et al., 2016) and are negative for Flock House virus infection. For insulin priming experiments, culture media was supplemented with 1.7 μM insulin from bovine pancreas (Sigma 10516), as used previously (Zhang et al., 2011). For ERK inhibition experiments, culture media was supplemented with 10 μM U0126 in DMSO (Cell Signaling 9903) (Pakpour et al., 2012; Surachetpong et al., 2009) 24 hours prior to and during infection.

West Nile virus-Kunjin (strain MRM16) was provided by R. Tesh, passaged twice in Vero cells, and purified by ultracentrifugation. Zika virus (Paraiba strain) was gifted by S. Best, passaged once in C6/36 cells, and purified by ultracentrifugation. Dengue virus 2 (New Guinea C strain) was gifted by S. Best. All experiments with a specific virus type utilized the same stock.

#### Mosquito rearing

*Culex quinquefasciatus* eggs were originally collected near Johannesburg, South Africa and distributed by the Centers for Disease Control and Prevention by BEI Resources, NIAID, NIH (*Cx. quinquefasciatus*, Strain JHB, Eggs, NR-43025), and mosquitoes were reared following the conditions described by BEI and following (Kauffman et al., 2017). Adult female mosquitoes were fed on defibrinated chicken blood (Colorado Serum Company 31141) in hog sausage casing. Adults were provided continuous access to sucrose (JT Baker 4072-01). 6-9 day old adult female mosquitoes were deprived of sucrose 48 hours prior to experimental feedings, as described (Moudy et al., 2009).

### METHOD DETAILS

#### Plaque assay

All data showing titers of KUNV, ZIKV, and DENV were determined by standard plaque assay on Vero cells (Baer and Kehn-Hall, 2014), with the exception of viral RNA levels in mosquitoes, which were determined by qRT-PCR.

#### Immunoprecipitation and Immunoblotting

Protein extracts were prepared by lysing cells with RIPA buffer (25 mM Tris-HCl (pH 7.6), 150 mM NaCl, 1 mM EDTA, 1% NP-40, 1% sodium deoxycholate, 0.1% SDS, 1mM Na3VO4, 1 mM NaF, 0.1 mM PMSF, 10 μM aprotinin, 5 μg/mL leupeptin, 1 μg/mL pepstatin A). Protein samples were diluted using 2x Laemmli loading buffer, mixed, and boiled for 5 minutes at 95 °C. Samples were analyzed by SDS/PAGE using a 10% acrylamide gel, followed by transfer onto PVDF membranes (Millipore IPVH00010). Membranes were blocked with 5% BSA (ThermoFisher BP9706) in Tris-buffered saline (50 mM Tris-HCl pH 7.5, 150 mM NaCl) and 0.1% Tween-20 for 1 hour at room temperature.

Following insulin treatment and virus infection in *D. melanogaster* S2 cells, cells were lysed in RIPA buffer and 100 μg of total protein were immunoprecipitated with an antibody recognizing total Stat92E (Santa Cruz, sc-15708) at 4 °C overnight, followed by a 2 hour incubation with Protein A agarose beads (Pierce 22811) at 4°C. Beads were washed three times with NETN buffer (20 mM Tris-HCl, pH 8.0, 100 mM NaCl, 1 mM EDTA, and 0.5% NP-40). Beads were boiled for 10 minutes at 95 °C and supernatants were subjected to Western blotting for P-Tyr (Cell Signaling 8954).

Primary antibody labeling was done with anti-P-Akt (Ser473) (1:2,000) (Cell Signaling 4060), anti-Akt (pan) (1:1,000) (Cell Signaling 4691), ERK (Cell Signaling 4695) (1:1,000), P-ERK (Cell Signaling 4370) (1:2,000), P-Tyr (Cell Signaling 8954S) (1:2,000), or anti-actin (Sigma A2066) (1:10,000) overnight at 4 °C. Secondary antibody labeling was done using anti-rabbit (Promega 4011) or anti-goat (Jackson Immunoresearch 705-035-147) IgG-HRP conjugate (1:10,000) by incubating membranes for 2 hours at room temperature. Blots were imaged onto film using luminol enhancer (ThermoFisher 1862124).

#### Quantitative reverse transcriptase PCR

qRT-PCR was used to measure gene mRNA levels in S2 cells and *Cx. quinquefasciatus*. Cells or mosquitoes were lysed with Trizol Reagent (ThermoFisher 15596). RNA was isolated by column purification (ZymoResearch R2050), DNA was removed (ThermoFisher 18068), and cDNA was prepared (BioRad 170–8891). Expression of *D. melanogaster* genes *InR*, *AGO1*, *AGO2*, *Dicer-2*, *upd2*, *upd3*, and *vir-1* were measured using SYBR Green reagents (ThermoFisher K0222) and normalized to *Rp49*. Expression of *TotM* was measured using primer/probe sets for *TotM* (Dm02362087_s1 ThermoFisher 4351372) and normalized to *RpL32* (Dm02151827_g1 (ThermoFisher 4331182), as in (Kemp et al., 2013), using TaqMan Universal Master Mix (ThermoFisher 4304437). Expression of the *Cx. quinquefasciatus* genes *CxR2D2* and *CxSTAT* and of the WNV-Kun *envelope* and *NS5* genes was measured using SYBR Green and normalized to *Cx18S*. The reaction for both SYBR Green and TaqMan samples included one cycle of denaturation at 95 °C for 10 minutes, followed by 40 cycles of denaturation at 95 °C for 15 seconds and extension at 60 °C for 1 minute, using an Applied Biosystems 7500 Fast Real Time PCR System. ROX was used as an internal control. qRT-PCR primer sequences are listed in the Resources table (Deddouche et al., 2008; Lin et al., 2004; Paradkar et al., 2012; Reid et al., 2015; Spellberg and Marr, 2015).

#### Larval infections and confocal microscopy

Third-instar *D. melanogaster* larvae were washed with PBS, placed in a pool of buffer with WNV-Kun, and pricked with a tungsten needle (Hiroyasu et al., 2018). Fat bodies were dissected, fixed, and blocked following (Loza-Coll et al., 2014). Samples were stained with DAPI, mounted onto coverslips using ProLong Diamond Antifade Mountant (Invitrogen P36961), and imaged using a Leica Sp8X confocal microscope.

#### Fly infections

2-7 day old adult *D. melanogaster* were anesthetized with CO2 and injected intrathoracically with 23 nL of WNV-Kun to achieve a dose of 10,000 PFU/fly, as described (Martin et al., 2018). Mock infection was the equivalent of injection with saline. For mortality studies, groups of at least 40 flies were injected and kept in vials containing cornmeal food (Genesee Scientific #66-112). DGRP survival curves were repeated to verify reproducibility. All survival studies for a specific mutant (e.g. *InR* RNAi or deletion mutants) were repeated and the survival data was combined. Vials were changed every three days. For viral titer experiments, three groups of five flies were collected, homogenized in PBS, and used as individual samples for plaque assay.

#### Mosquito infections

WNV-Kun was added to chicken blood washed with RPMI, supplemented with 20% FBS and HEPES buffer or 170 pM insulin (Kang et al., 2008; Pakpour et al., 2012), at a final concentration of 1×107 PFU WNV-Kun/mL. Female *Cx. quinquefasciatus* were maintained for three generations and subsequently fed using an artificial mosquito feeder (Chemglass Life Sciences) for 2 hours. Engorged females were separated under CO2, placed in 1 gal cartons with continuous access to 10% sucrose, and collected at 1, 5, and 10 days post-infection for analysis. Females that did not feed were excluded from all subsequent analysis.

### QUANTIFICATION AND STATISTICAL ANALYSIS

Results presented as dot plots show data from individual biological replicates and the arithmetic mean of the data, shown as a black horizontal line. Biological replicates of adult flies consisted of five pooled animals and replicates of mosquitoes consisted of three pooled animals. Results shown are representative of at least duplicate experiments. All statistical analyses were completed using GraphPad Prism. Two-tailed unpaired t-tests assuming unequal variance were utilized to compare normally distributed pair-wise quantitative data. Two-way ANOVA with Tukey’s correction for multiple comparisons was used to compare multivariate data. Mann-Whitney tests were utilized to compare distribution-free quantitative data. All error bars represent standard deviation of the mean. Any statistical outliers for experiments were identified using a Grubb’s test (α=0.05) and removed. Survival curves were analyzed by the log-rank (Mantel-Cox) test using GraphPad Prism to determine P-values between infected genotypes.

### DATA AND SOFTWARE AVAILABILITY

All input phenotypic data (hazard ratios) are provided in Table S1. Full output GWAS data is available upon request.

### SUPPLEMENTAL INFORMATION

**Table S1.** Phenotype input data for genome-widess association study.

| DGRP ID# | Log(hazard ratio) | DGRP ID# | Log(hazard ratio) | DGRP ID# | Log(hazard ratio) |
| --- | --- | --- | --- | --- | --- |
| 26 | 0.349471799 | 358 | 0.471878199 | 810 | 0.801746619 |
| 31 | 0.355259906 | 359 | 0.104828404 | 812 | 0.72542155 |
| 32 | 0.34869419 | 367 | 0.102776615 | 843 | 0.790988475 |
| 38 | 0.183269844 | 373 | 0.050379756 | 849 | -0.055319966 |
| 41 | 0.40534636 | 375 | 0.260309946 | 857 | 0.187520721 |
| 42 | 0.165244326 | 377 | -0.003007018 | 894 | 0.895864351 |
| 45 | 0.062957834 | 379 | 0.404320467 | 900 | 0.229937686 |
| 57 | -0.089428952 | 385 | 0.050766311 | 907 | 0.471144965 |
| 59 | 0.292034436 | 386 | -0.066057397 | 908 | -0.059035507 |
| 83 | -0.13288622 | 390 | 0.116607744 | 911 | 0.035429738 |
| 85 | 0.216693599 | 391 | 0.615002615 |  |  |
| 88 | 0.046885191 | 392 | 0.551327988 |  |  |
| 91 | 0.004321374 | 395 | 0.494154594 |  |  |
| 101 | 0.003029471 | 399 | 0.805364907 |  |  |
| 105 | -0.021317435 | 406 | 0.520614522 |  |  |
| 129 | 0.079904468 | 426 | 0.36604921 |  |  |
| 138 | 0.396896449 | 427 | 0.614580867 |  |  |
| 158 | 0.23350376 | 437 | 0.630834518 |  |  |
| 161 | -0.041483897 | 439 | 0.42553422 |  |  |
| 177 | 0.267640982 | 443 | 0.716504164 |  |  |
| 195 | 0.14176323 | 491 | -0.323763783 |  |  |
| 208 | 0.059941888 | 492 | 0.333245699 |  |  |
| 217 | 0.151982395 | 502 | 1.083860801 |  |  |
| 228 | 0.127104798 | 508 | 0.312388949 |  |  |
| 229 | 0.946992341 | 509 | 0.710286648 |  |  |
| 235 | 0.083860801 | 517 | 0.133538908 |  |  |
| 239 | 0.302979937 | 559 | 0.639386869 |  |  |
| 301 | 0.371621927 | 563 | 1.051152522 |  |  |
| 303 | 0.753123245 | 566 | 0.325515663 |  |  |
| 307 | 0.464340485 | 596 | -0.137510833 |  |  |
| 309 | 0.072249898 | 627 | 0.471731651 |  |  |
| 313 | 0.176958981 | 703 | 0.294686624 |  |  |
| 315 | 0.415474168 | 714 | 0.934952708 |  |  |
| 324 | 0.120902818 | 732 | -0.078261516 |  |  |
| 332 | 0.237794993 | 757 | 0.143951116 |  |  |
| 348 | 0.352182518 | 765 | 0.256958153 |  |  |
| 350 | 0.021602716 | 774 | 0.053078443 |  |  |
| 354 | 0.555336328 | 799 | 0.725993259 |  |  |
| 357 | 0.168497484 | 808 | 0.281487888 |  |  |

**Table S2.**
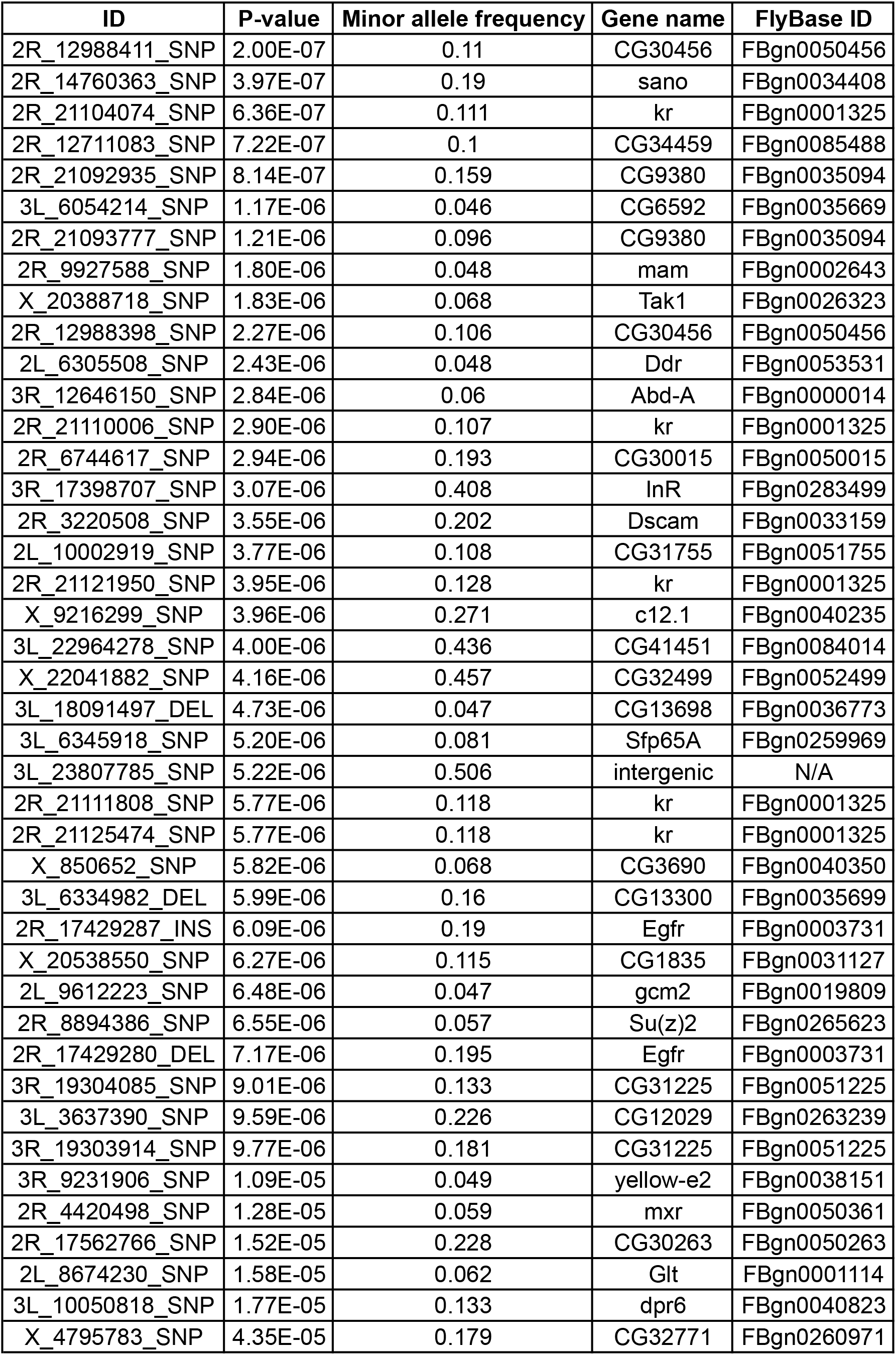
List of genome-wide suggestive variants (P < 5E-05)

| ID | P-value | Minor allele frequency | Gene name | FlyBase ID |
| --- | --- | --- | --- | --- |
| 2R_12988411_SNP | 2.00E-07 | 0.11 | CG30456 | FBgn0050456 |
| 2R_14760363_SNP | 3.97E-07 | 0.19 | sano | FBgn0034408 |
| 2R_21104074_SNP | 6.36E-07 | 0.111 | kr | FBgn0001325 |
| 2R_12711083_SNP | 7.22E-07 | 0.1 | CG34459 | FBgn0085488 |
| 2R_21092935_SNP | 8.14E-07 | 0.159 | CG9380 | FBgn0035094 |
| 3L_6054214_SNP | 1.17E-06 | 0.046 | CG6592 | FBgn0035669 |
| 2R_21093777_SNP | 1.21E-06 | 0.096 | CG9380 | FBgn0035094 |
| 2R_9927588_SNP | 1.80E-06 | 0.048 | mam | FBgn0002643 |
| X_20388718_SNP | 1.83E-06 | 0.068 | Tak1 | FBgn0026323 |
| 2R_12988398_SNP | 2.27E-06 | 0.106 | CG30456 | FBgn0050456 |
| 2L_6305508_SNP | 2.43E-06 | 0.048 | Ddr | FBgn0053531 |
| 3R_12646150_SNP | 2.84E-06 | 0.06 | Abd-A | FBgn0000014 |
| 2R_21110006_SNP | 2.90E-06 | 0.107 | kr | FBgn0001325 |
| 2R_6744617_SNP | 2.94E-06 | 0.193 | CG30015 | FBgn0050015 |
| 3R_17398707_SNP | 3.07E-06 | 0.408 | InR | FBgn0283499 |
| 2R_3220508_SNP | 3.55E-06 | 0.202 | Dscam | FBgn0033159 |
| 2L_10002919_SNP | 3.77E-06 | 0.108 | CG31755 | FBgn0051755 |
| 2R_21121950_SNP | 3.95E-06 | 0.128 | kr | FBgn0001325 |
| X_9216299_SNP | 3.96E-06 | 0.271 | c12.1 | FBgn0040235 |
| 3L_22964278_SNP | 4.00E-06 | 0.436 | CG41451 | FBgn0084014 |
| X_22041882_SNP | 4.16E-06 | 0.457 | CG32499 | FBgn0052499 |
| 3L_18091497_DEL | 4.73E-06 | 0.047 | CG13698 | FBgn0036773 |
| 3L_6345918_SNP | 5.20E-06 | 0.081 | Sfp65A | FBgn0259969 |
| 3L_23807785_SNP | 5.22E-06 | 0.506 | intergenic | N/A |
| 2R_21111808_SNP | 5.77E-06 | 0.118 | kr | FBgn0001325 |
| 2R_21125474_SNP | 5.77E-06 | 0.118 | kr | FBgn0001325 |
| X_850652_SNP | 5.82E-06 | 0.068 | CG3690 | FBgn0040350 |
| 3L_6334982_DEL | 5.99E-06 | 0.16 | CG13300 | FBgn0035699 |
| 2R_17429287_INS | 6.09E-06 | 0.19 | Egfr | FBgn0003731 |
| X_20538550_SNP | 6.27E-06 | 0.115 | CG1835 | FBgn0031127 |
| 2L_9612223_SNP | 6.48E-06 | 0.047 | gcm2 | FBgn0019809 |
| 2R_8894386_SNP | 6.55E-06 | 0.057 | Su(z)2 | FBgn0265623 |
| 2R_17429280_DEL | 7.17E-06 | 0.195 | Egfr | FBgn0003731 |
| 3R_19304085_SNP | 9.01E-06 | 0.133 | CG31225 | FBgn0051225 |
| 3L_3637390_SNP | 9.59E-06 | 0.226 | CG12029 | FBgn0263239 |
| 3R_19303914_SNP | 9.77E-06 | 0.181 | CG31225 | FBgn0051225 |
| 3R_9231906_SNP | 1.09E-05 | 0.049 | yellow-e2 | FBgn0038151 |
| 2R_4420498_SNP | 1.28E-05 | 0.059 | mxr | FBgn0050361 |
| 2R_17562766_SNP | 1.52E-05 | 0.228 | CG30263 | FBgn0050263 |
| 2L_8674230_SNP | 1.58E-05 | 0.062 | Glt | FBgn0001114 |
| 3L_10050818_SNP | 1.77E-05 | 0.133 | dpr6 | FBgn0040823 |
| X_4795783_SNP | 4.35E-05 | 0.179 | CG32771 | FBgn0260971 |

**Table S3.**
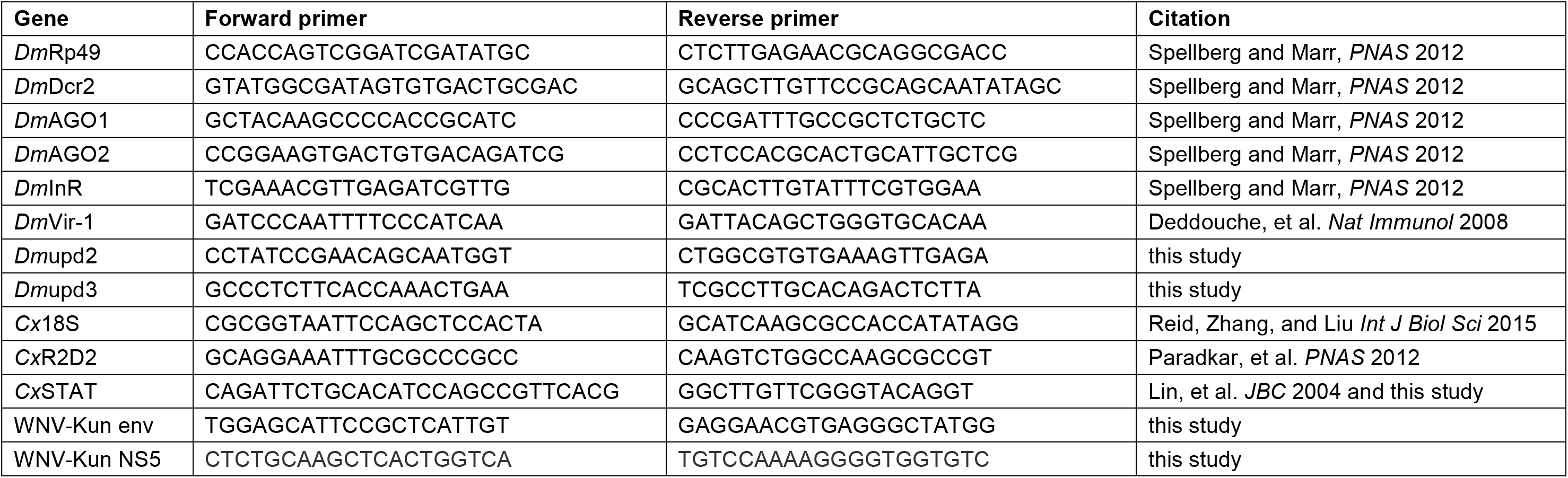
qRT-PCR Primers.

**Fig. S1:**
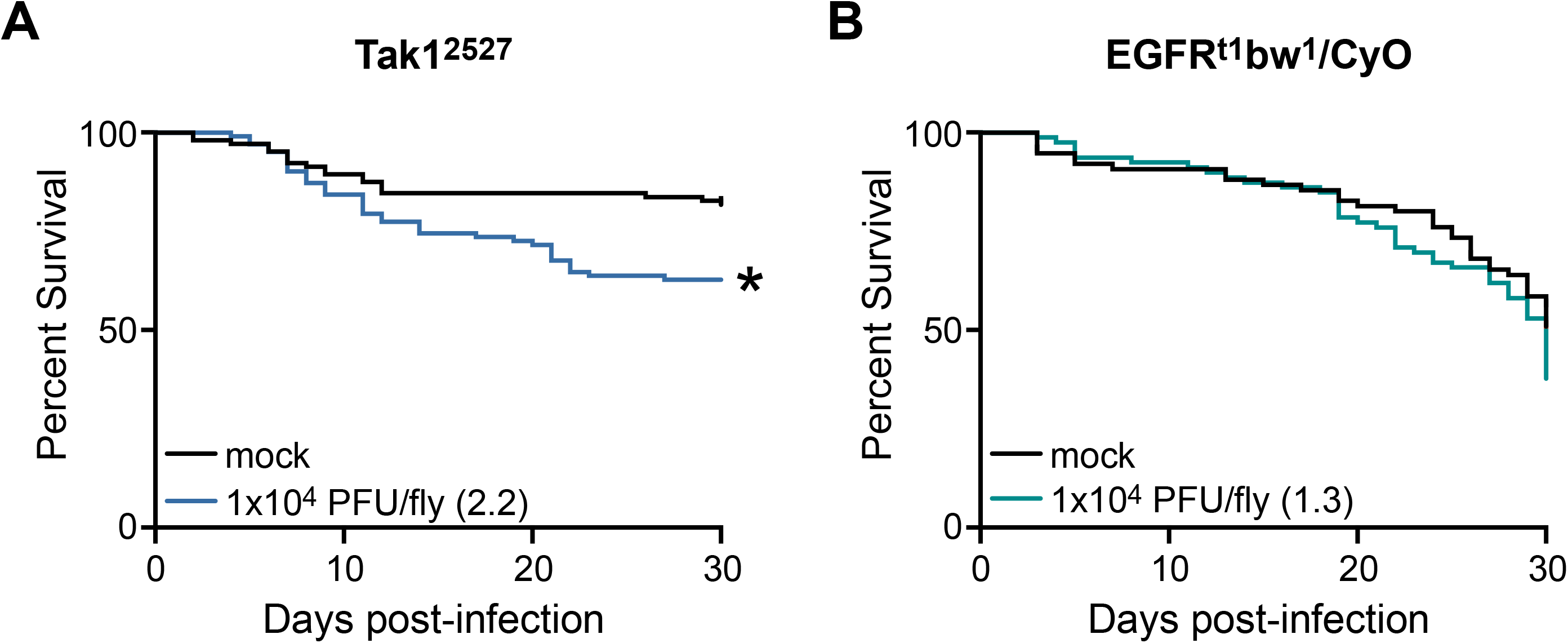
Validation of candidate genes identified by GWAS analysis. (A) *Tak1* and (B) *EGFR*-mutant flies were infected with WNV-Kun and survival was monitored for 30 days post-infection. Hazard ratio for each infection group is indicated in parenthesis and statistical significance from the mock infection group is indicated with an asterisk. (Log-rank test; *p < 0.05). Each survival curve represents two independent experiments of >40 flies that were combined for a final survival curve.

**Fig. S2:**
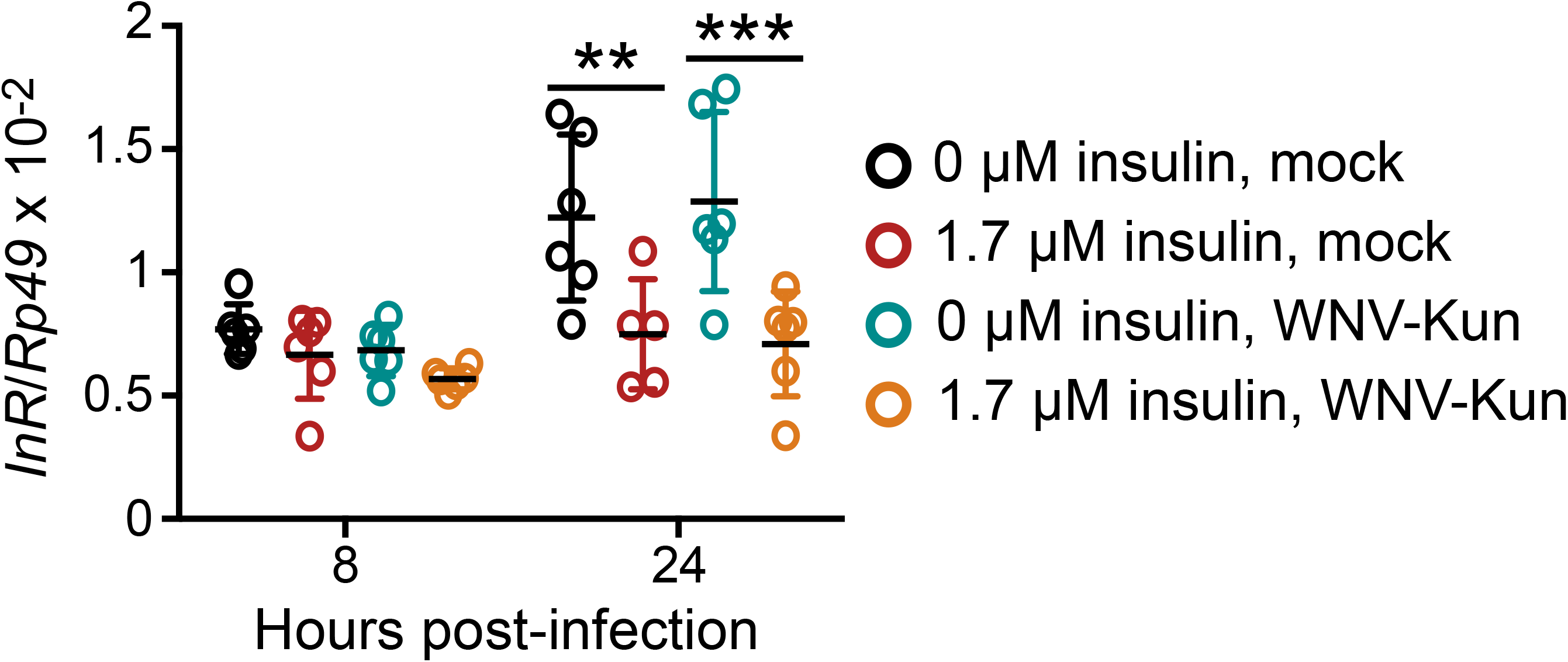
Insulin-treated S2 cells have reduced insulin receptor expression. Induction of *InR* was measured by qRT-PCR following priming of *Drosophila* S2 cells with insulin and mock or WNV-Kun infection. (**p < 0.01; ***p < 0.001). Error bars represent SDs.

**Fig. S3:**
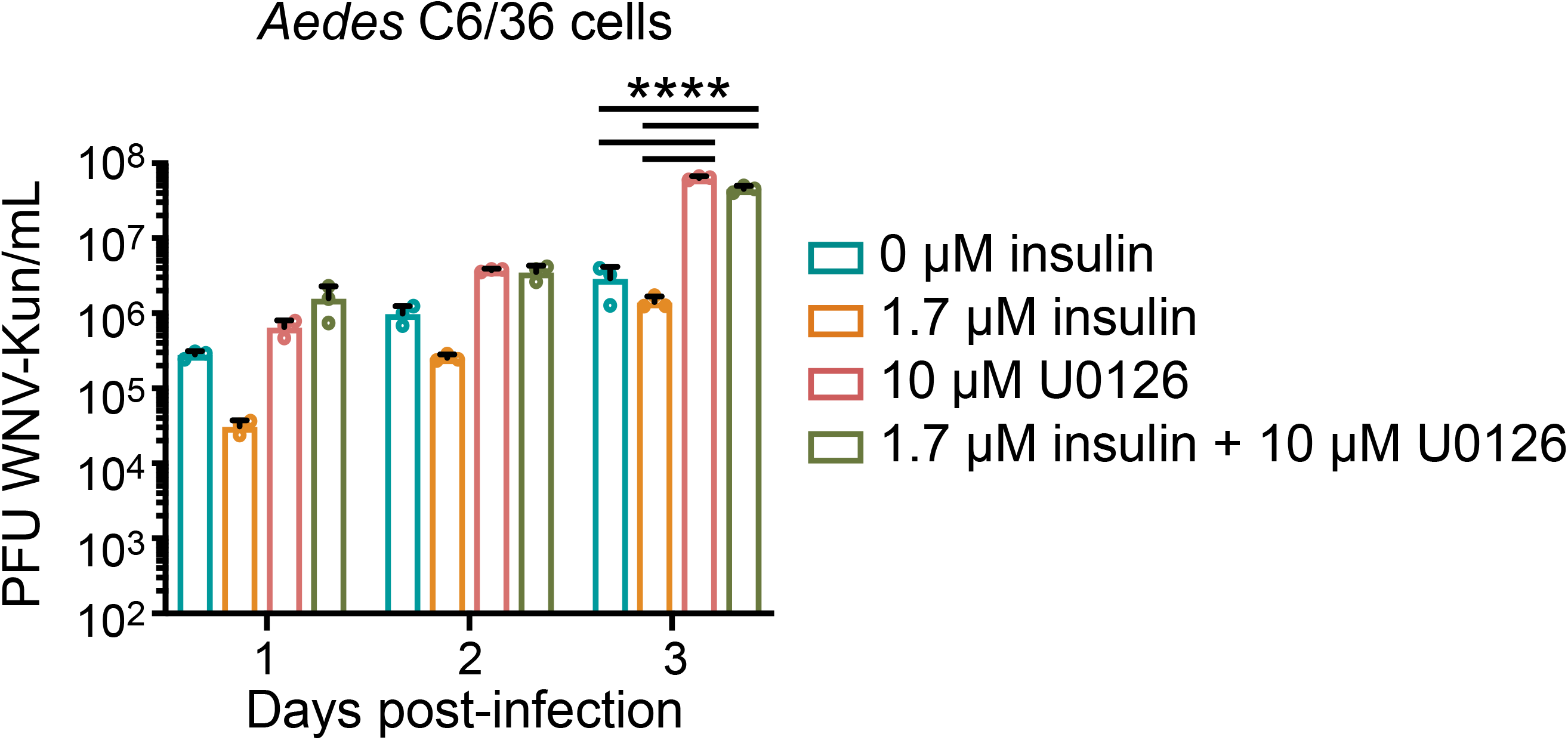
WNV-Kun titer is increased in ERK-inhibited cells. *Aedes* C6/36 cells were primed with 1.7 µM insulin, 10 µM U0126, or both for 24 hours prior to infection with WNV-Kun (MOI 0.01). Supernatant was collected and viral titer was measured by plaque assay. (****p < 0.0001). Error bars represent SDs.

## Resources Table

**Table S1.**
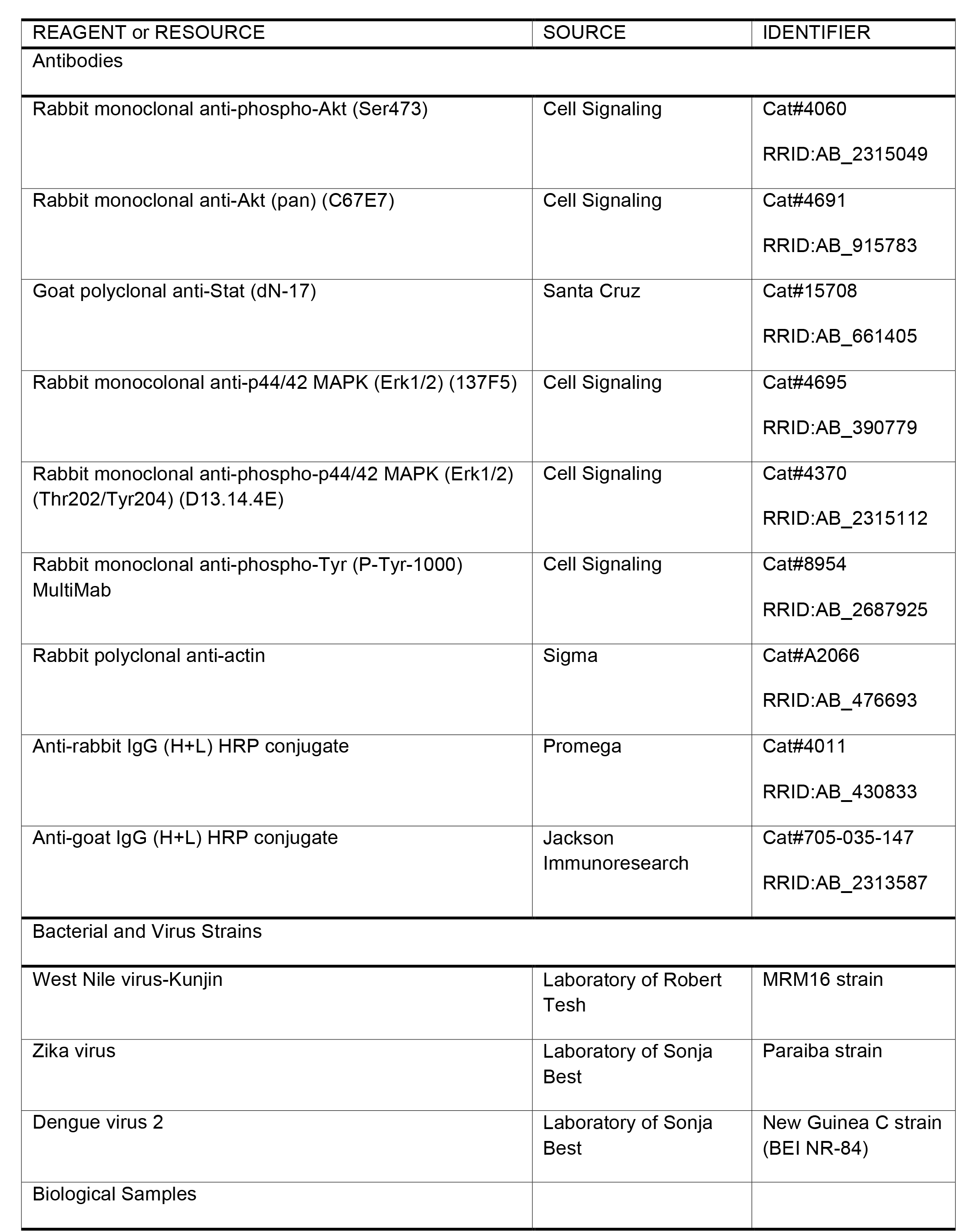
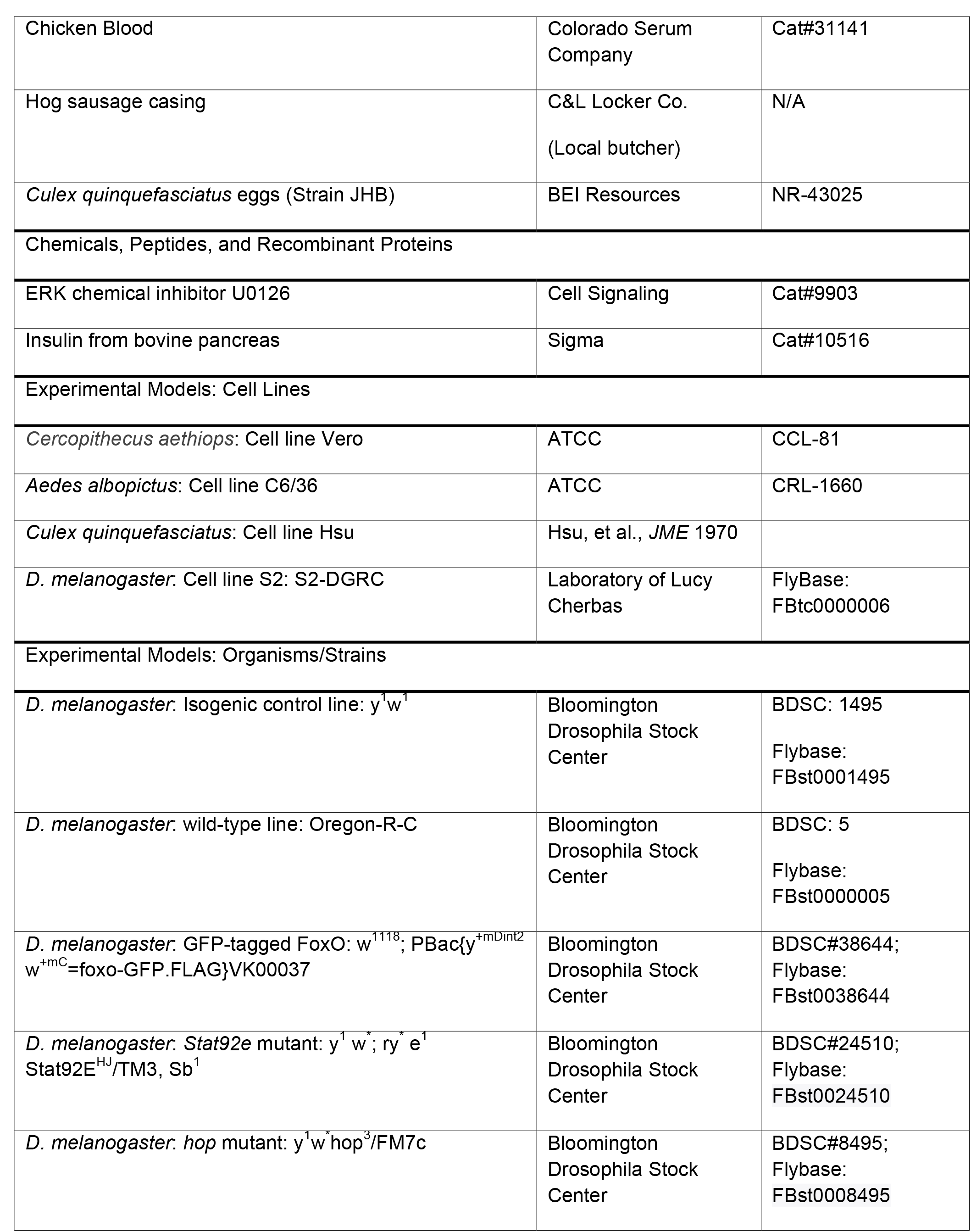
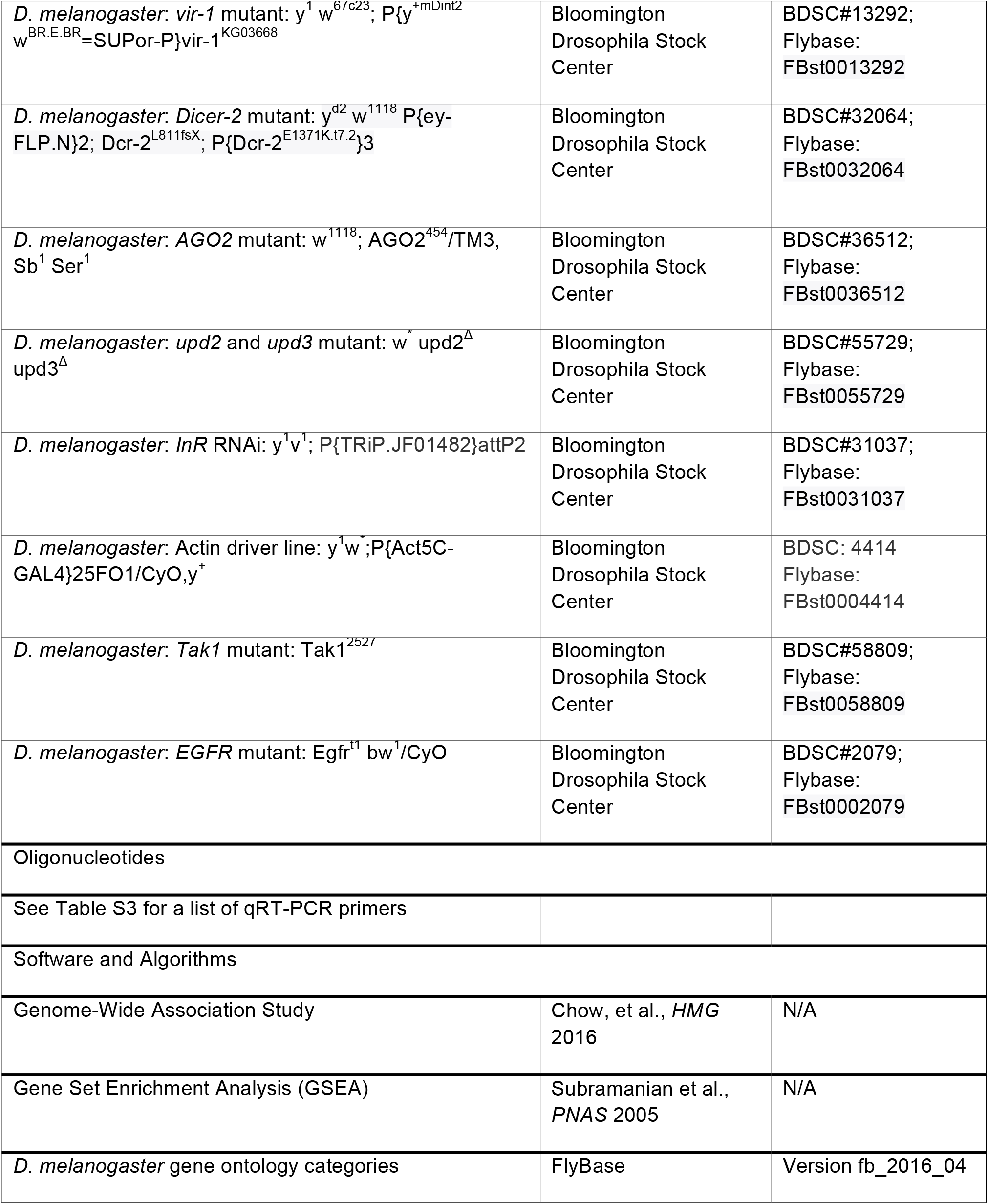
Phenotype input data for genome-widess association study.

